## Supplementary material for "Side chain to main chain hydrogen bonds stabilize a polyglutamine helix in the activation domain of a transcription factor": SI

Supplementary figures

Figure S1: Prediction of helical and Ncap propensity, for peptides $uQ_{25}$, $uL_4Q_{25}$ and $L_4Q_{25}$ obtained by using Agadir [1]. The peptides have been aligned so that the Gln residues, that are common to all of them, have the same residue numbers (Q11-Q35).

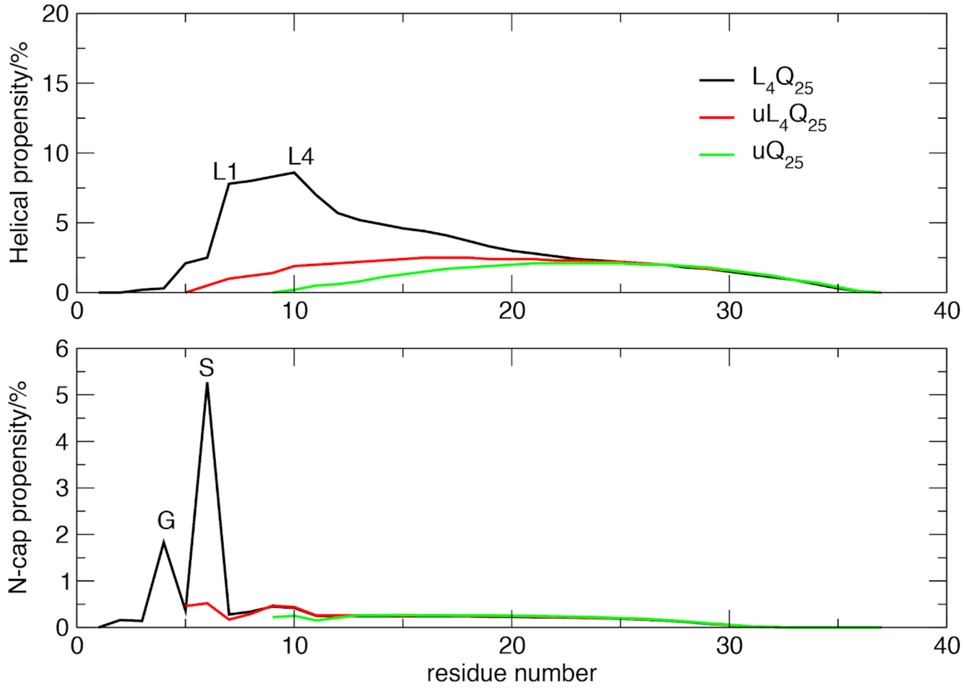

Figure S2: CD spectra of $uQ_{25}$, $uL_4Q_{25}$ and $L_4Q_{25}$.

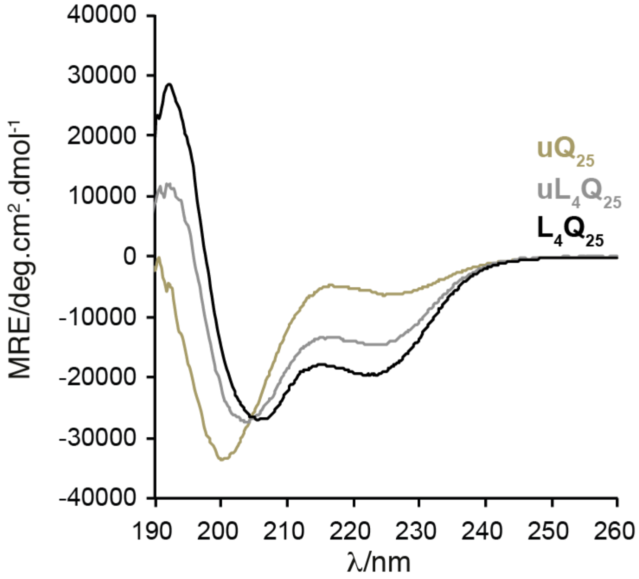

Figure S3: Region of the $^{1}H,^{15}N$ HSQC spectrum of peptides $L_4Q_4$ to $L_4Q_{16}$ showing the resonances belonging to the backbone amide groups of Gln residues.

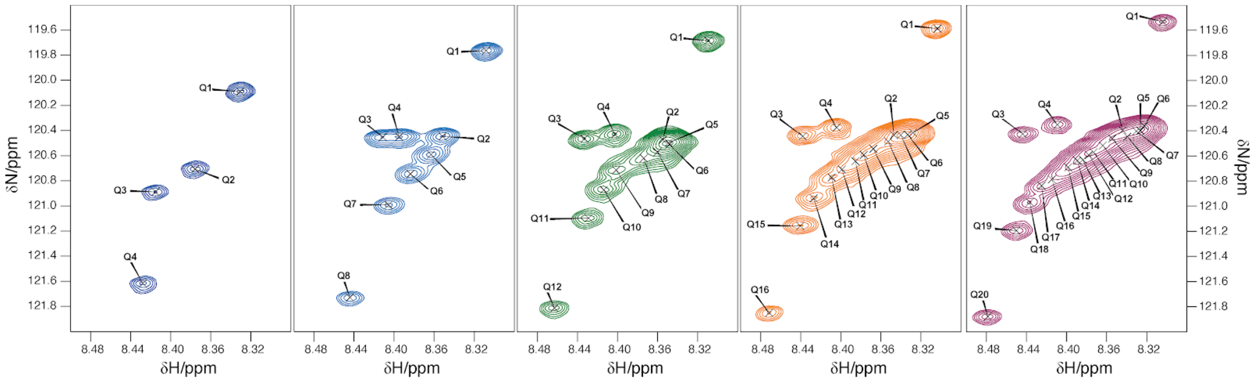

Figure S4: HNCO (light) and HN(CA)CO (dark) spectra of peptides $L_4Q_4$ to $L_4Q_{16}$ used for the assignment of the backbone resonances.

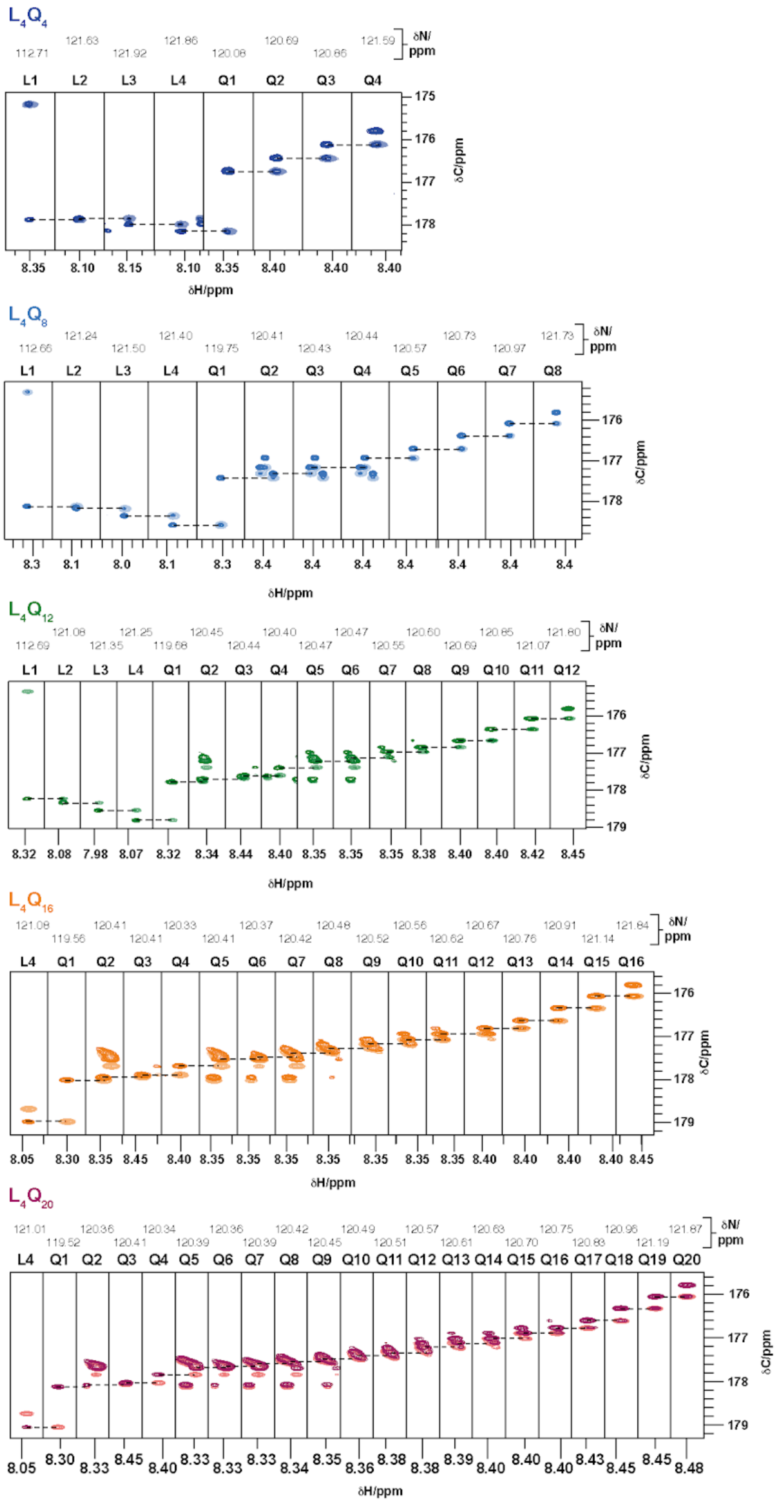

Figure S5: Strip plots showing the Cα cross-peaks correlated to the main chain NH resonances in the HN(CO)CACB experiment (light) and side chain NH$\varepsilon_{21}$ resonances in the (H)CC(CO)NH experiment (dark), used in the assignment of the latter.

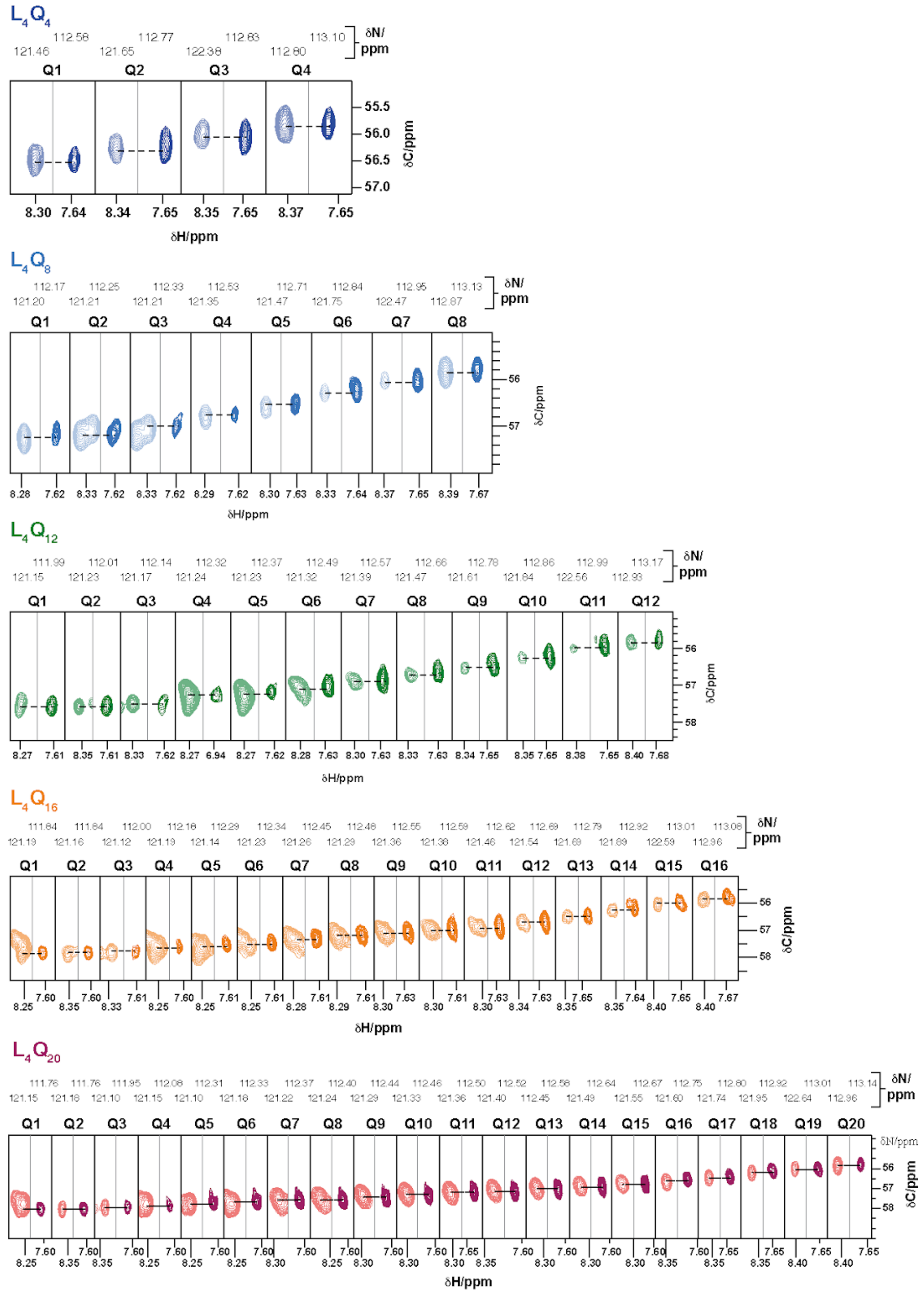

Figure S6: H$\varepsilon_{21}$-centered regions of the $^{15}$N planes of the H(CC)(CO)NH spectra of peptides $L_4Q_4$ to $L_4Q_{20}$ containing the side chain aliphatic $^1$H resonances of all residues of the polyQ tract.

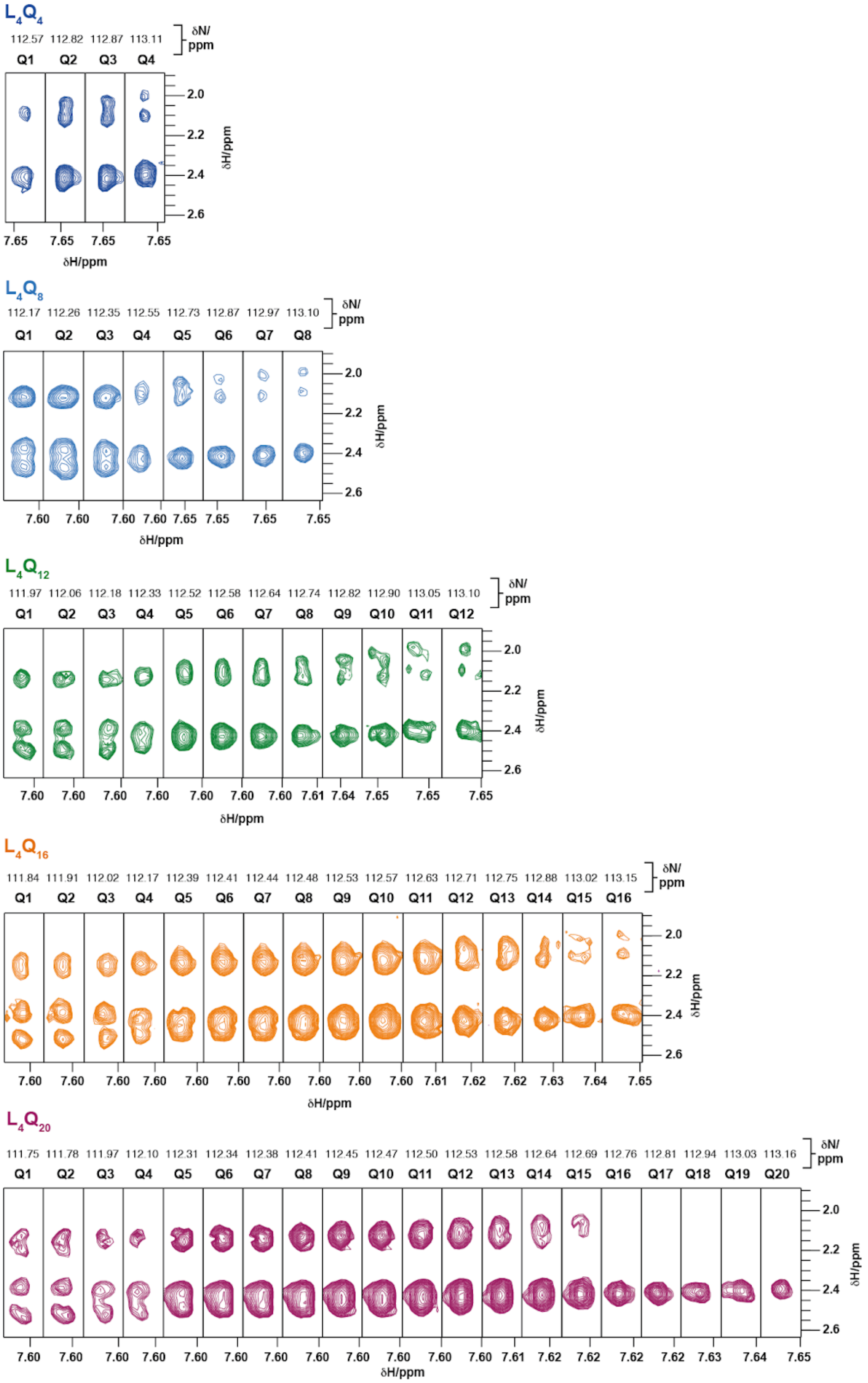

Figure S7: Time series of the secondary structure of peptides $L_4Q_4$ to $L_4Q_{20}$ as obtained by using the algorithm DSSP [2] where H stands for helix, in red; E stands for extended, in dark gray; and C stands for coil, in light gray.

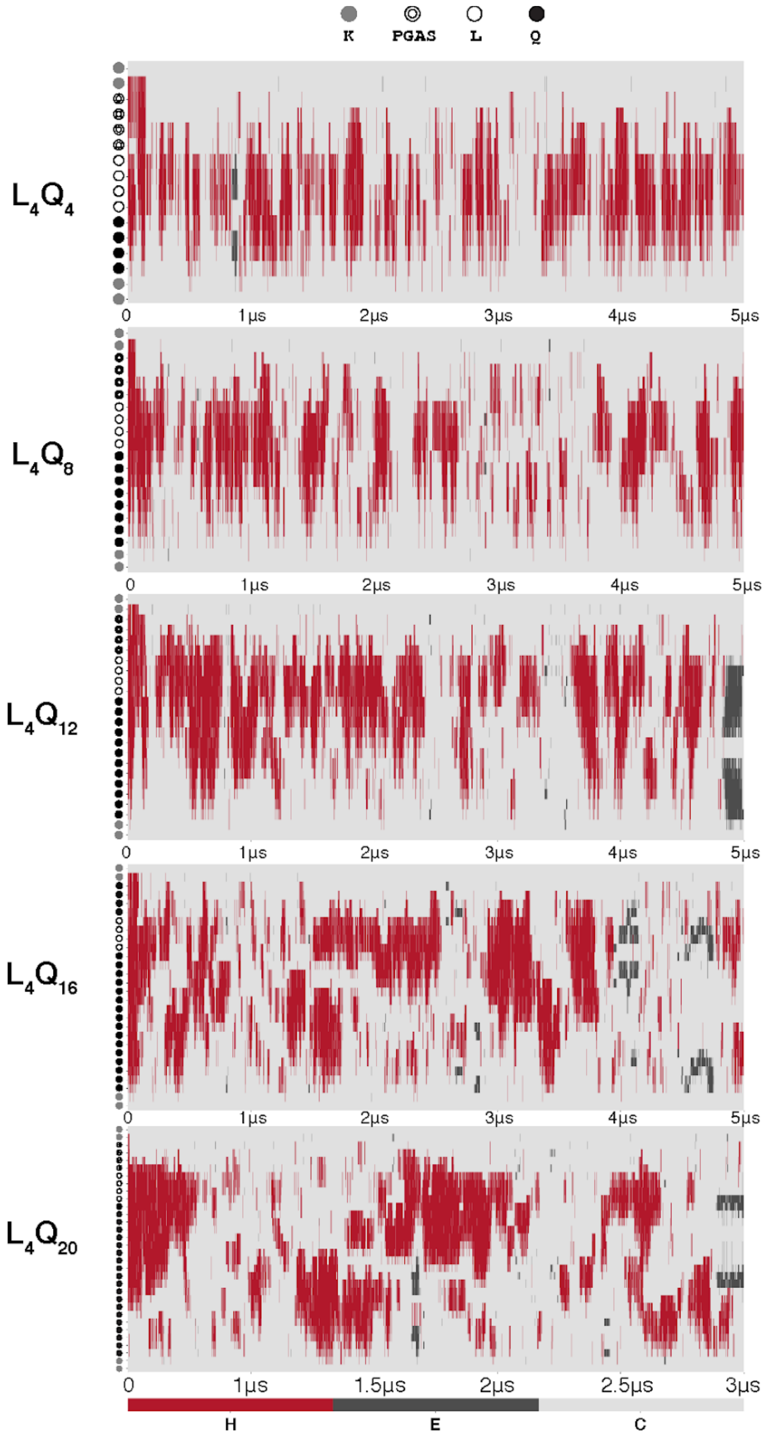

Figure S8: Experimental and back-calculated (a) Cα and (b) C' chemical shifts of peptides $L_4Q_4$ to $L_4Q_{20}$ from the reweighted trajectories obtained with different values of the parameter θ, that determines the degree of reweighting applied in the ME algorithm.

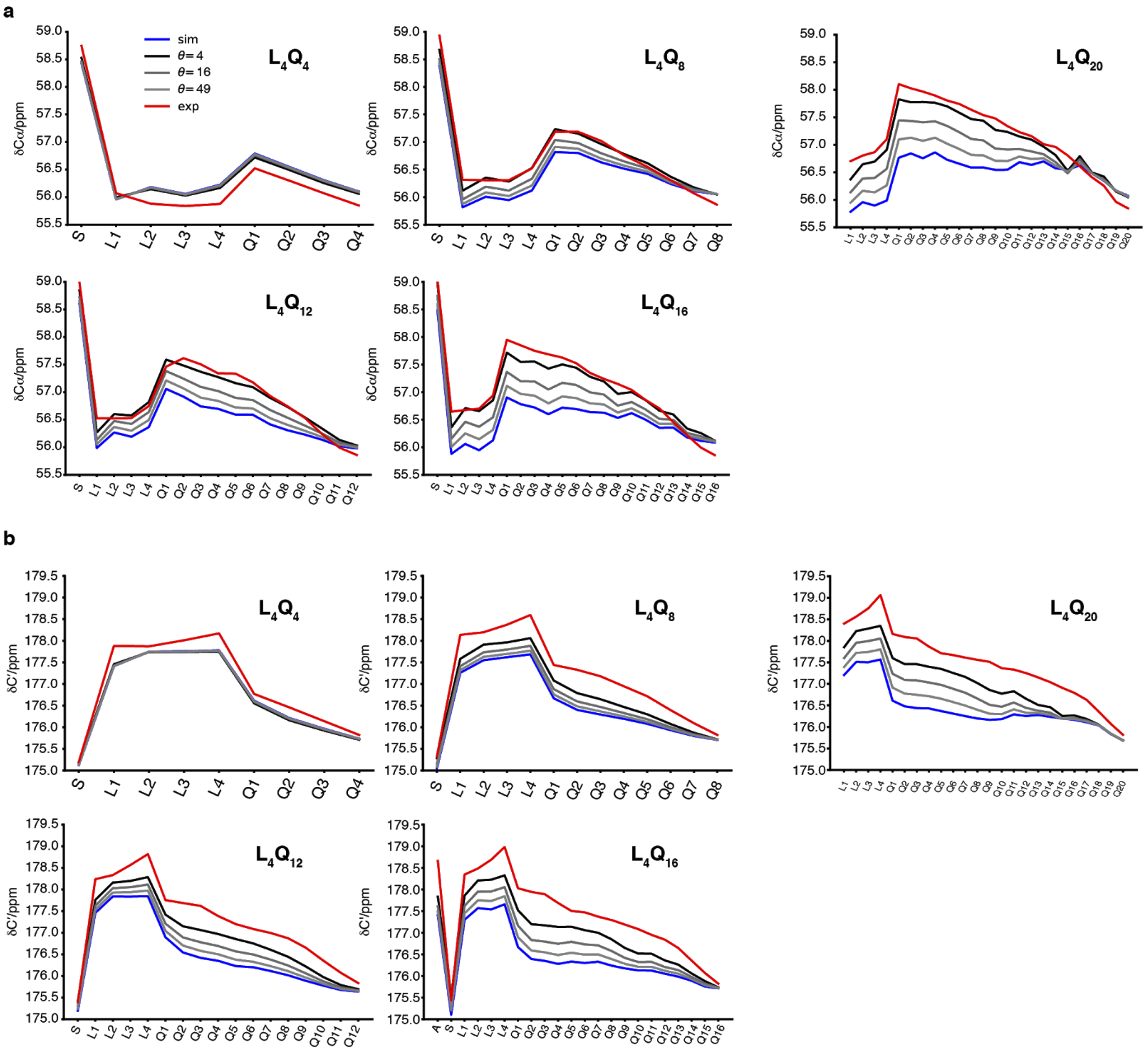

Figure S9: Effective fraction of frames after reweighting vs $\chi^2$ for different values of θ. $N_{eff}$ was calculated as $\exp(S_{rel})$ with $S_{rel}$ being the relative entropy term in the BME reweighting approach. In short, $N_{eff}$ quantifies the effective fraction of frames that are left after the trajectory has been reweighted to fit the data to an extent measured by $\chi^2$.

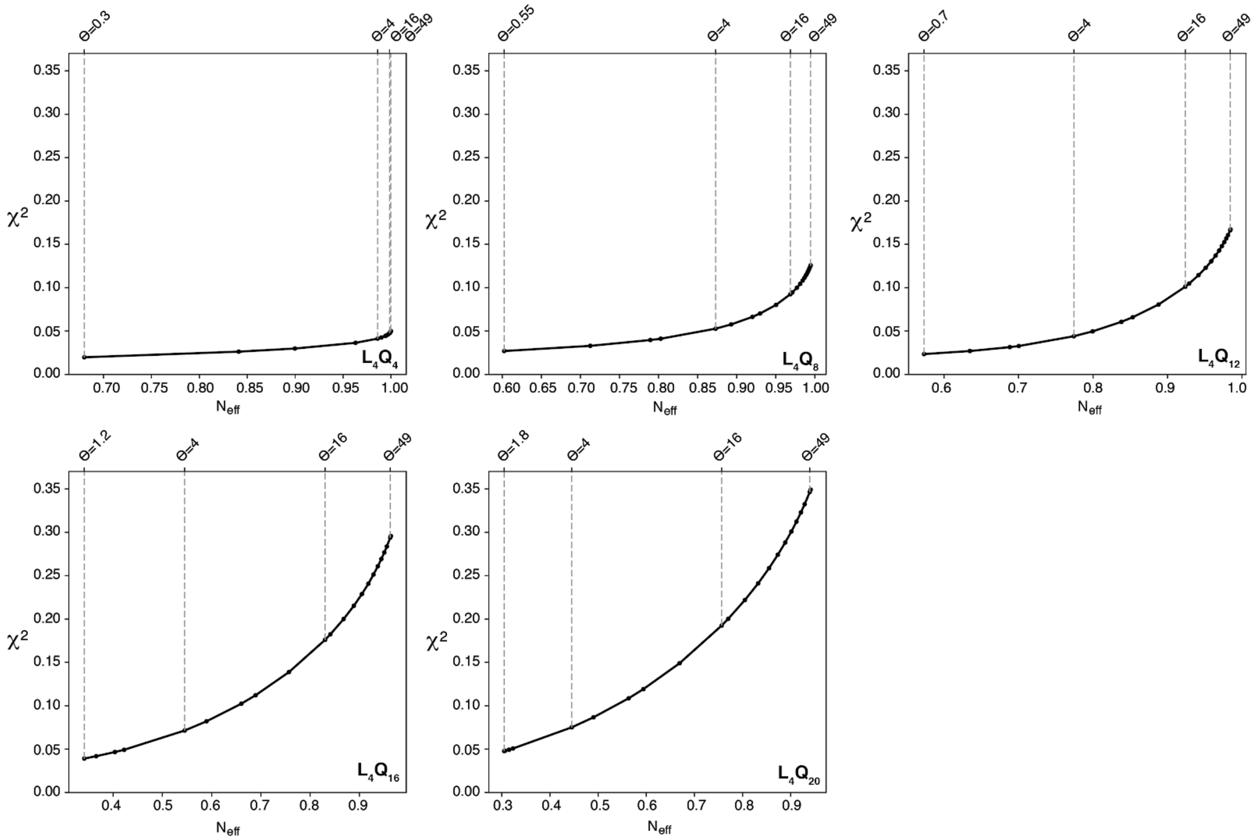

Figure S10: 2D histogram, in red, of backbone torsion angles φ and ψ for segments of five residues where the first and last residue are involved in a (i+4→i) side chain to main chain hydrogen bond. In black we show the result obtained when these residues are not involved in such interaction.

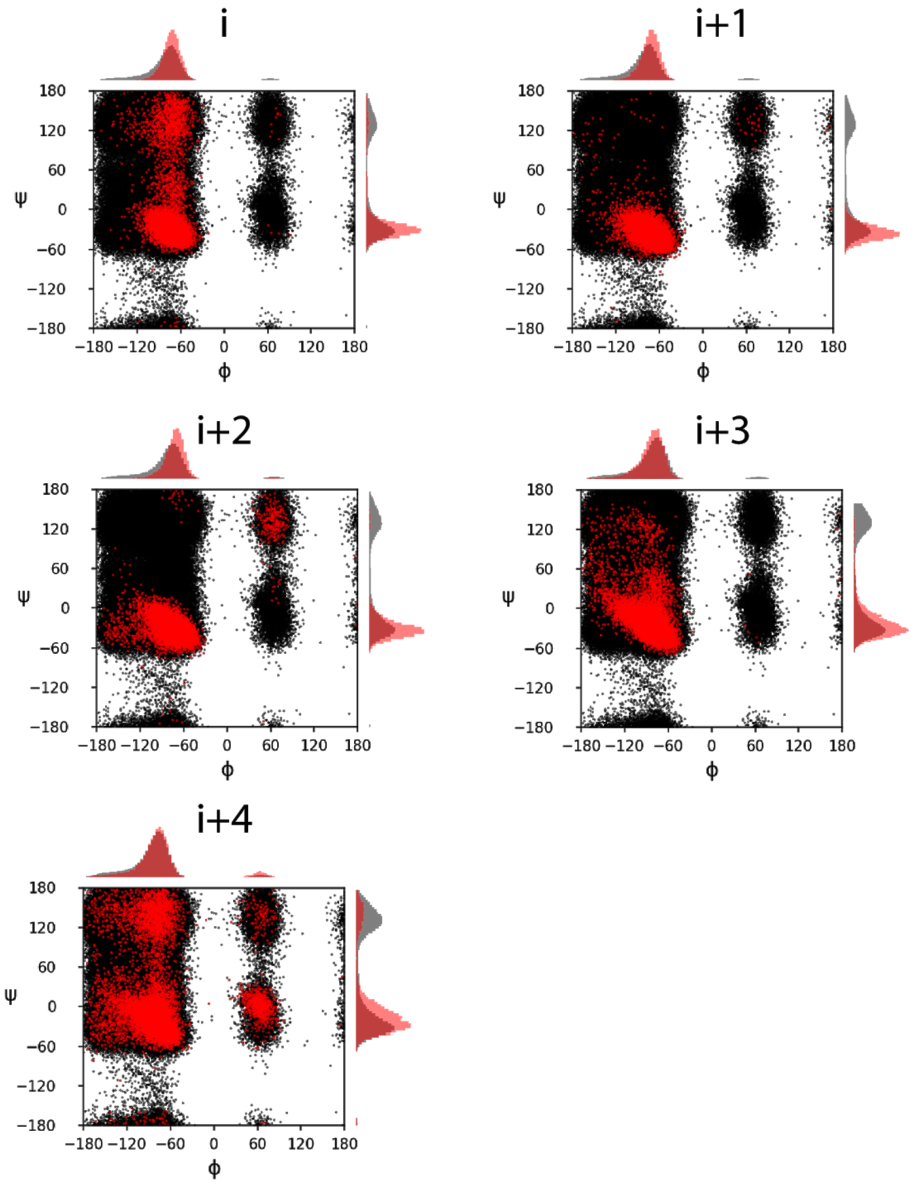

Figure S11: Comparison of the CD spectra of peptides Q1E to Q5E, as well as that of peptide Q1-4E, to that of peptide $L_4Q_{16}$ , shown in orange in all plots.

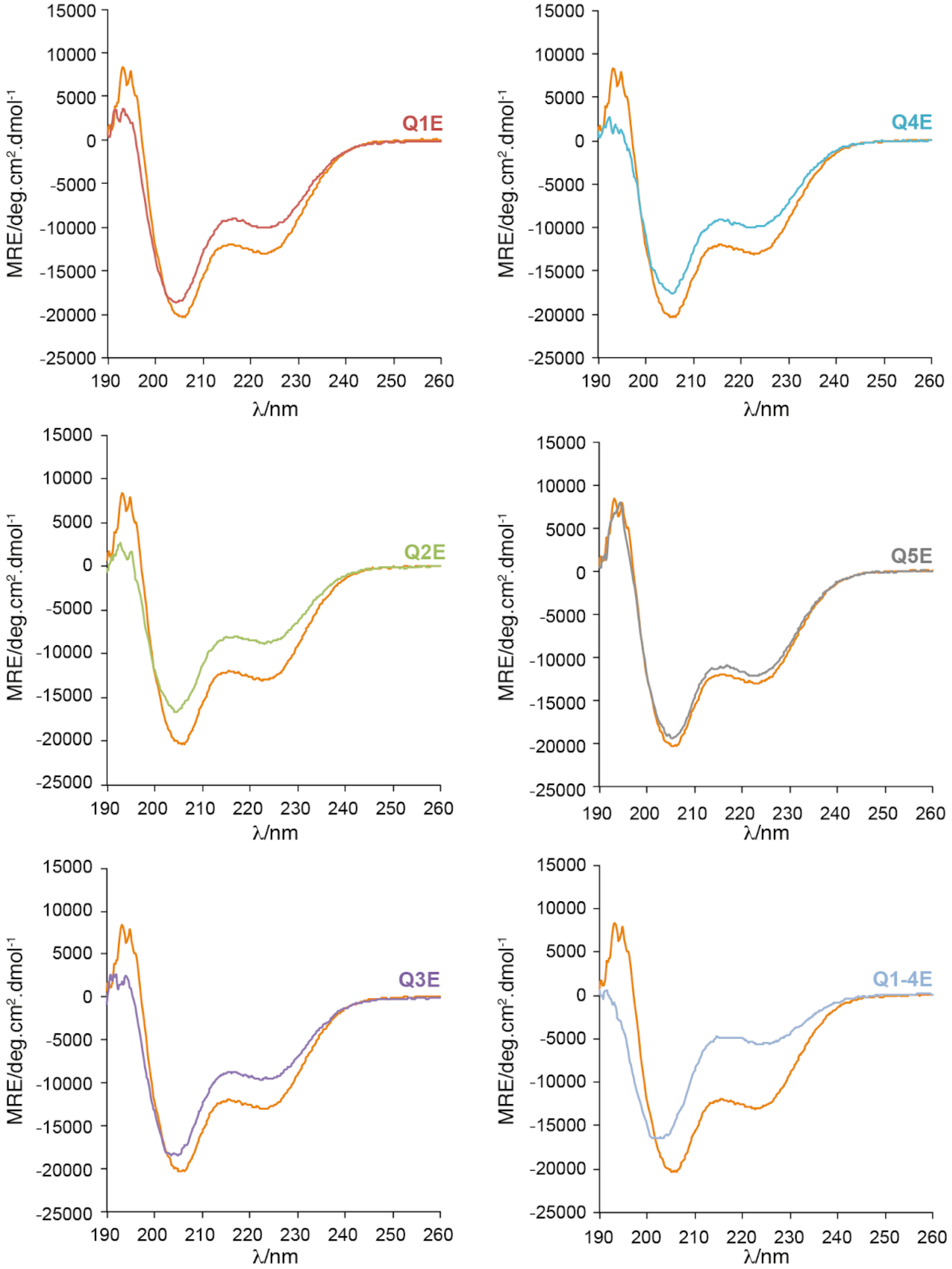

Figure S12: Comparison of the CD spectrum of peptide L1-4A, to that of peptide $L_4Q_{16}$.

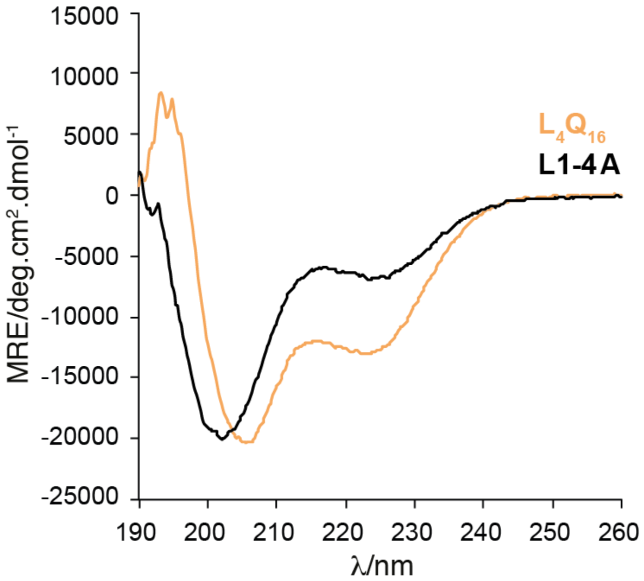

Figure S13: Prediction of the helicities of peptides $L_4Q_{16}$ and L1-4A obtained by using Agadir [1].

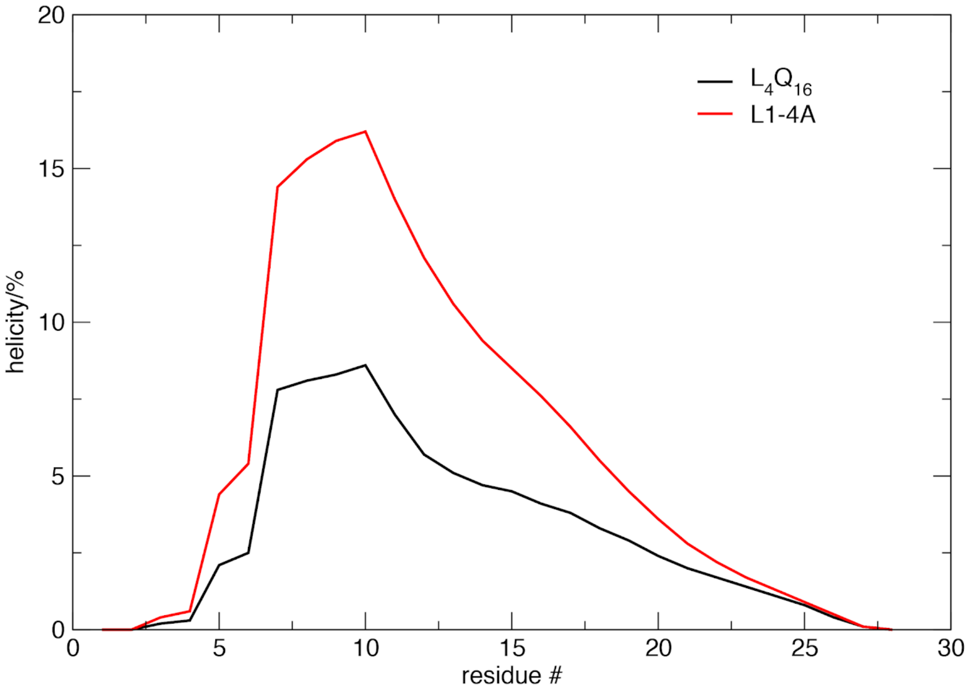

Figure S14: CD spectra of peptides $L_4Q_{16}$ (in orange) and Q1-4E at pH 2

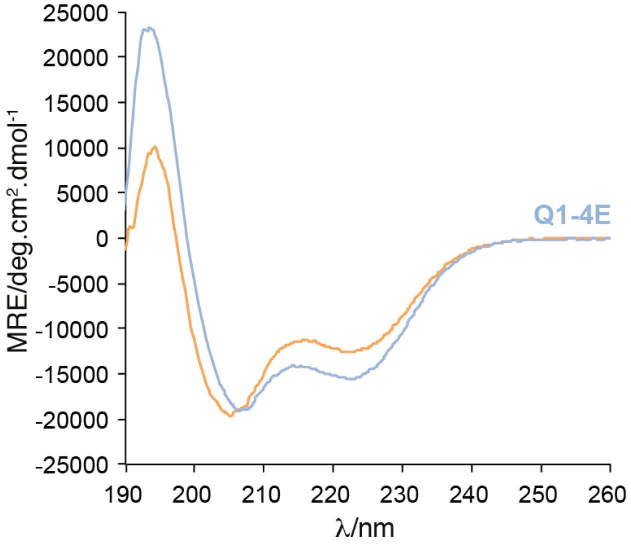

Figure S15: Distribution of the distance between the main chain NH of Q1 and the main chain CO of L1 in the absence and in the presence of the sc(Q1)→mc(L1) hydrogen bond.

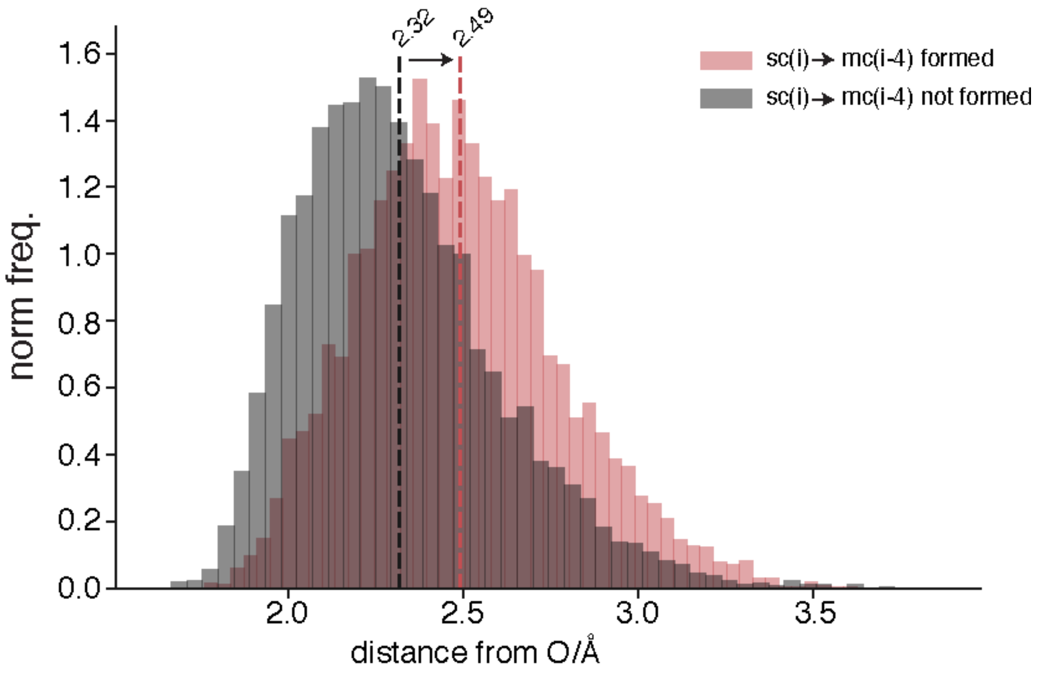
